# Low-cost open hardware system for behavioural experiments simultaneously with electrophysiological recordings

**DOI:** 10.1101/821843

**Authors:** Leandro A. A. Aguiar, Nivaldo A P de Vasconcelos, Gabriela Chiuffa Tunes, Antonio J. Fontenele, Romildo de Albuquerque Nogueira, Marcelo Bussotti Reyes, Pedro V. Carelli

## Abstract

A major frontier in neuroscience is to find neural correlates of perception, learning, decision making, and a variety of other types of behavior. In the last decades, modern devices allow simultaneous recordings of different operant responses and the electrical activity of large neuronal populations. However, the commercially available instruments for studying operant conditioning are expensive, and the design of low-cost chambers has emerged as an appealing alternative to resource-limited laboratories engaged in animal behavior. In this article, we provide a full description of a platform that records the operant behavior and synchronizes it with the electrophysiological activity. The programming of this platform is open source, flexible and adaptable to a wide range of operant conditioning tasks. We also show results of operant conditioning experiments with freely moving rats with simultaneous electrophysiological recordings.

**Specifications table:** 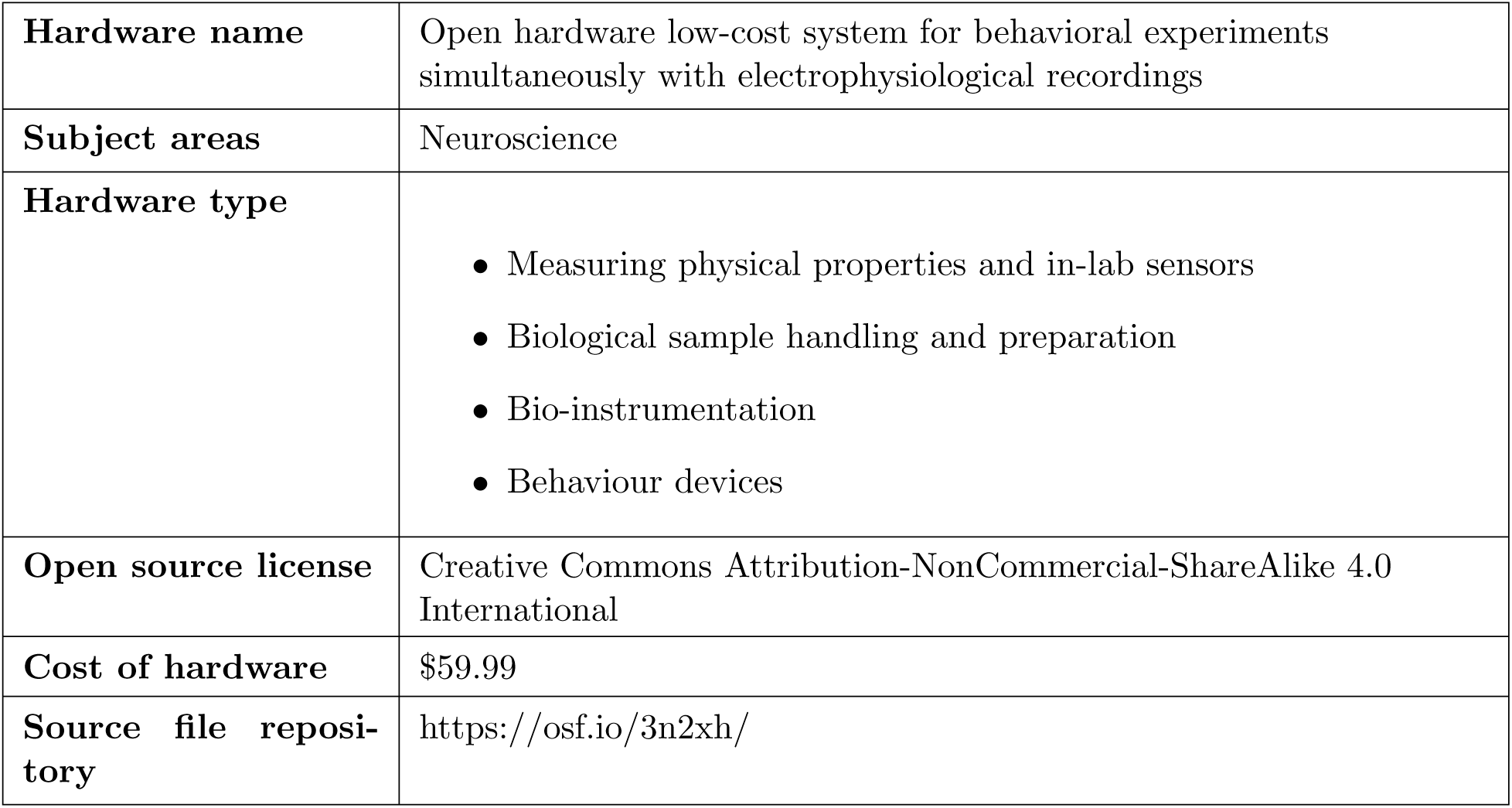

## 1. Hardware in context

Understanding the neural substrate of behavioral processes is a primary aim in modern neuroscience [1]. A central approach to study the neural substrate of behavior consists of identifying fluctuations of the neuronal activity associated with specific behavioral states – the neural correlates of behavior [2, 3, 4, 5]. However, neuronal activity and operant behavior have widely different hardware requirements. Therefore, experimenters usually use distinct recording systems for different kinds of data. This diversity of data-recording systems requires a mechanism of synchronization between them.

Neuronal activity originates from the transmembrane currents in the extracellular medium [6]. The electrical potential in the extracellular medium can be measured by using an implanted electrode in the target brain area. In the 1990s, technological advances allowed the development of multi-electrode array (MEA) manufacturing [7, 8]. MEAs have enabled low-cost recording of the extracellular activity of large neuronal populations in the brain in awake animals, including freely moving animals [9, 10, 11, 12].

Introduced by Skinner [13], the operant conditioning chamber (OCC) provided a new form of studying behavior, establishing the foundation over which an entire field of research developed. Currently, OCC – also called Skinner box – is a widely used device to perform behavioral studies based on animal models [1].

Commercial OCC systems may be costly. This can be a problem to resource-limited researchers [14]. High-cost creates financial obstacles to the scientific development as a whole. Thus, the design of low-cost OCCs has emerged as an appealing alternative to resource-limited laboratories run behavioral studies based on animal models. Some research groups have developed low-cost OCC [14, 15]. However, none of them provided an efficient system able to send information to the electrophysiology recording system regarding the behavioral events produced in the OCC.

Arduino is a user friendly and very flexible open electronic prototyping platform family [16]. It is able to monitor and analyze input and output signals through digital and analog ports, as well as sending and receiving information through Transistor-Transistor Logic (TTL). It is based on microcontrollers (eg Microchip ATmega328P) fast enough to accurately describe behavioral or electrophysiological activity. Further, it has a serial communication port for communication with the computer via Universal Serial Bus (USB) [14]. This fine temporal resolution makes the Arduino a feasable low-cost device to synchronize behavioral and electrophysiological data [17].

In this paper, we developed a low-cost OCC, able to perform different behavioral tests, simultaneously sending synchronization pulses through an optic fiber to be recorded by the electrophysiological acquisition system. Thus both behavioral and electrophysiological data are stored together.

## 2. Hardware description

The Arduino-based systems have been successfully used to simultaneously record neural activity and behavior [18, 19]. Among the recent studies, the most similar system to ours is composed of a behavioral system integrated into two-photon imaging experiments, during a GoNoGo task [20]. In that system, they used water as reward, and the Arduino was responsible for controlling the water dispenser. However, the synchronization between the two-photon imaging, and the behavioral system was done by the serial port, which can be a source of noise for electrophysiological experiments. We use sugar pellets as a reward, released by a food pellet dispenser disk. It requires a more sophisticated releasing mechanism, when compared with water dispenser control [20, 21].

The main features of our system are:

- We developed and tested a low-cost behavioral system, based on Arduino microcontroller, which is able to synchronize behavioral experiments and electrophysiological recordings;
- We have used an optical link to relay the signal from the operant conditioning chamber to the acquisition system, avoiding noise contamination in the recordings.
- The system is also free hardware, free software and easily adaptable to different behavioral tasks.
- The main contribution of our systems is to provide an affordable and flexible system to investigate the neural correlates of behavior.

## 3. Design files

### 3.1 Design Files Summary

| Design filename | File type | Open source license | Location of the file |
| --- | --- | --- | --- |
| optical_receptor.JPG | image | CC BY-NC-SA 4.0 | Available with the article |
| optical_receptor_copper_bottom.gbr | pcb design | CC BY-NC-SA 4.0 | Available with the article |
| optical_receptor_silkscreen_bottom.gbr | pcb design | CC BY-NC-SA 4.0 | Available with the article |
| optical_receptor_silkscreen_top.gbr | pcb design | CC BY-NC-SA 4.0 | Available with the article |
| optical_receptor_soldermask_bottom.gbr | pcb design | CC BY-NC-SA 4.0 | Available with the article |
| optical_sender.JPG | image | CC BY-NC-SA 4.0 | Available with the article |
| optic_sender_copper_bottom.gbr | pcb design | CC BY-NC-SA 4.0 | Available with the article |
| optic_sender_silkscreen_bottom.gbr | pcb design | CC BY-NC-SA 4.0 | Available with the article |
| optic_sender_silkscreen_top.gbr | pcb design | CC BY-NC-SA 4.0 | Available with the article |
| optic_sender_soldermask_bottom.gbr | pcb design | CC BY-NC-SA 4.0 | Available with the article |
| disk.JPG | image | CC BY-NC-SA 4.0 | Available with the article |
| disk_copper_bottom.gbr | pcb design | CC BY-NC-SA 4.0 | Available with the article |
| disk_silkscreen_bottom.gbr | pcb design | CC BY-NC-SA 4.0 | Available with the article |
| disk_silkscreen_top.gbr | pcb design | CC BY-NC-SA 4.0 | Available with the article |
| disk_smdmask_bottom.gbr | pcb design | CC BY-NC-SA 4.0 | Available with the article |
| disk_soldermask_bottom.gbr | pcb design | CC BY-NC-SA 4.0 | Available with the article |
| box_all.pdf | pdf | CC BY-NC-SA 4.0 | Available with the article |
| disk.pdf | pdf | CC BY-NC-SA 4.0 | Available with the article |
| sideA.pdf,sideBB.pdf, sideCC.pdf and sideD.pdf | pdf | CC BY-NC-SA 4.0 | Available with the article |

## 4. Bill of materials

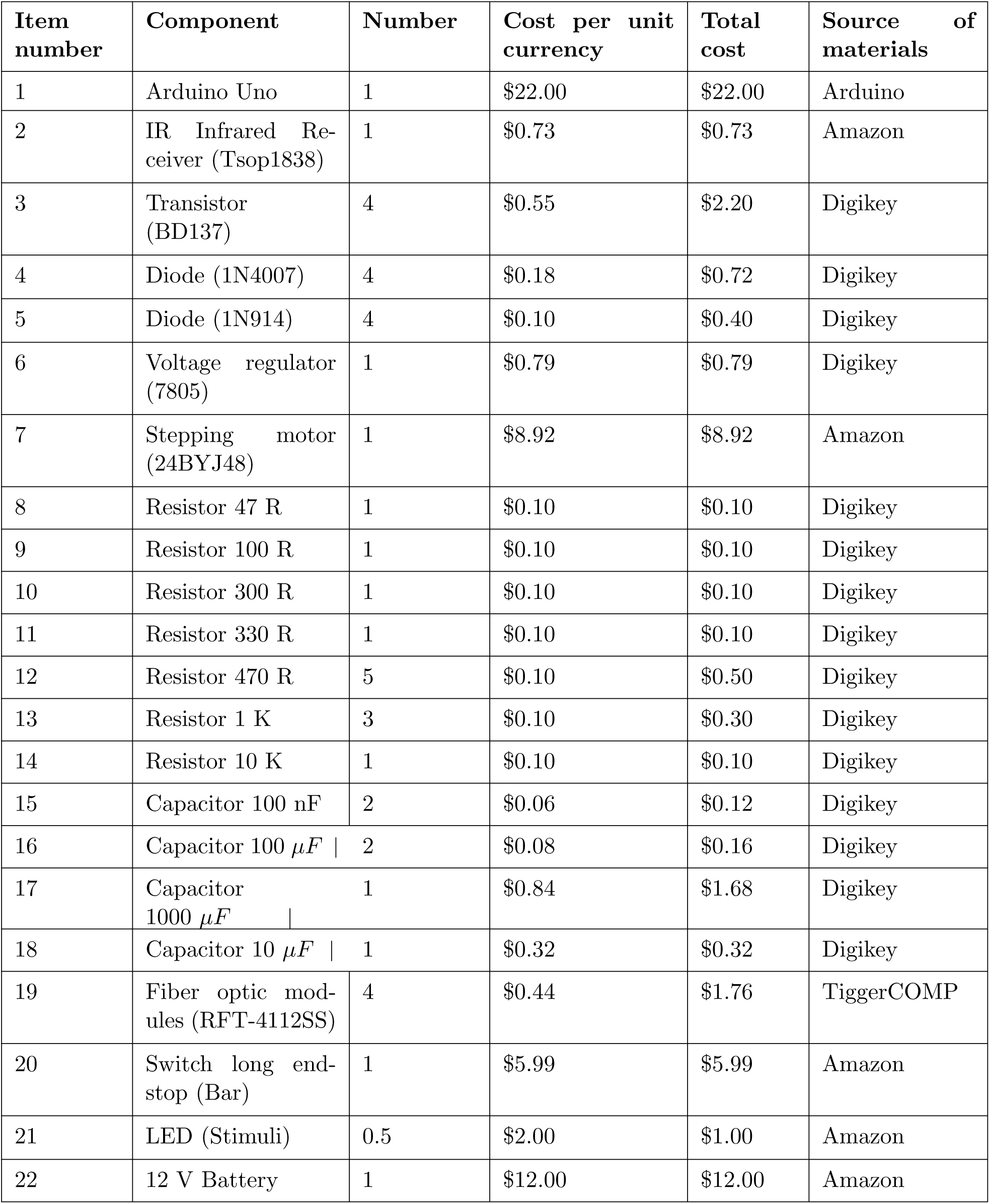

## 5. Build instructions

Our behavioral system can be seen in Figure 1a. The operant conditioning chamber consists of a cubicle measuring 25 *×* 25 *×* 30 cm, with walls based on 6 mm recycled acrylic plates (Fig. 1b). We used one of the lateral acrylic plate to attach the mechanical and electronic devices: food dispenser disk, an Arduino Uno microcontroller, an auxiliary electronic board, a stepper motor, and a device for optical coupling with the recording system (Fig. 1c). All mechanical and electronic materials were low-cost, easily purchased from electronics stores.

**Figure 1:**
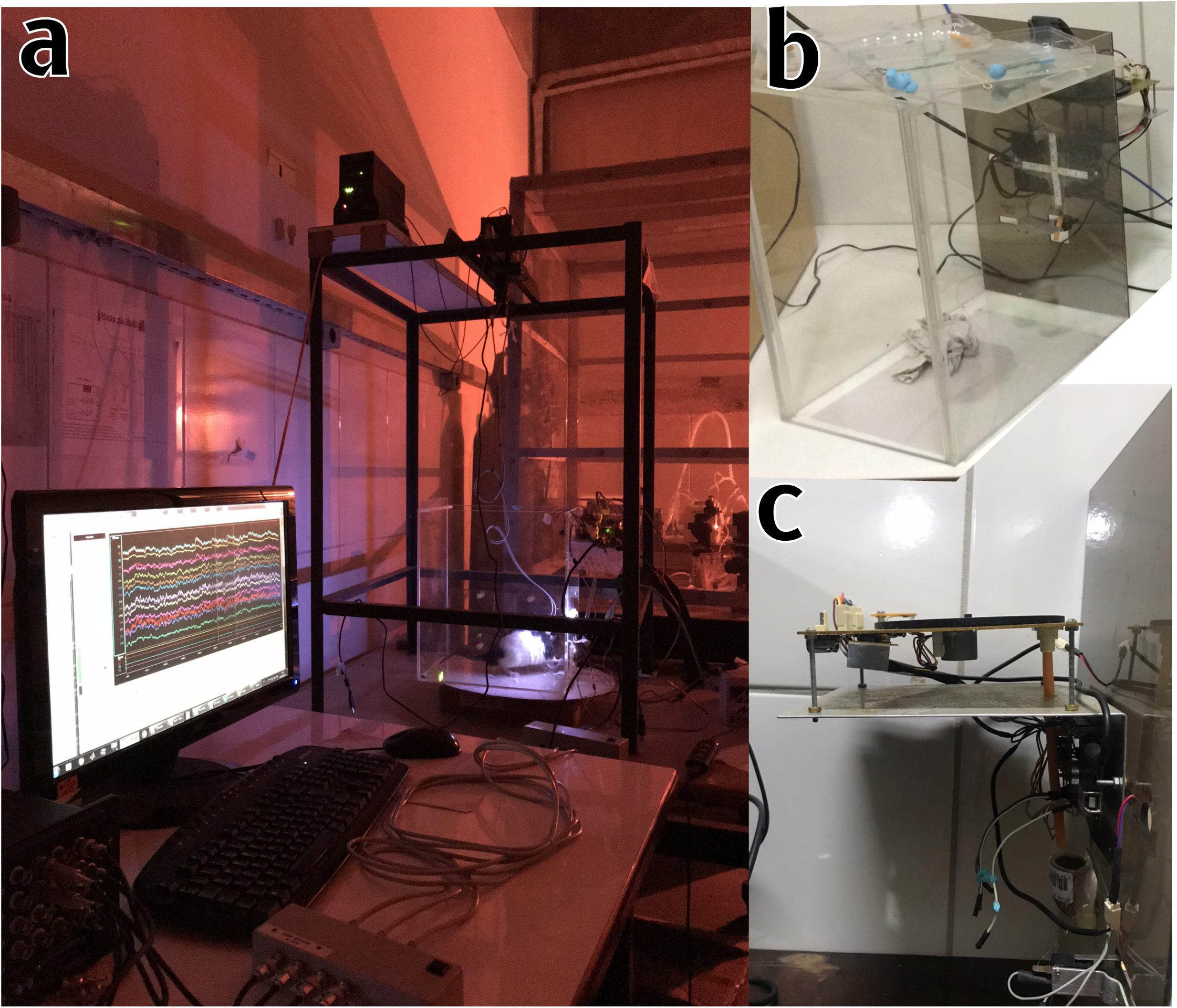
Picture of the complete system. **(a)** One animal in a training session in our system. It is possible to identify the rat – illuminated in the background – and the acrylic walls of the behavior box. **(b)** Closer view of the operant conditioning chamber. **(c)** Lateral view of the chamber zooming at the pellet dispenser and the electronic devices attached outside of the lateral wall.

The program run in the microcontroller must be able to autonomously control the entire behavioral experiment: monitoring the lever state (pressed or released), generating the sensory stimuli, evaluating the task contingencies, and delivering the reward. All controlling and informing signals were based on TTL pulses. An acrylic plate was used to build a chamber. The design files are freely available at the OSF repository [22].

### 5.1 Food pellet dispenser disk

We designed the food pellet dispenser disk to release 50 mg sugar pellets, which work as a reward in behavioral tasks (Fig.2). The dispenser disk was cut from a 4 mm polymethyl methacrylate (acrylic) plate and coupled to the stepper motor shaft. The disk has two groups of radially-distributed holes. One group consists of larger wholes that accommodate and drag the pellets until they are delivered (Fig. 2c - item 14). The other group has smaller holes (0.05 mm in diameter, Fig. 2c - item 15). They provide a mechanism of alignment of the disk rotation - when correctly aligned, the infrared light passes through the hole reaching the sensor, which in turn sends a TTL to the Arduino board (Fig. 2 a and c - items 5 and 6). This mechanism avoids cumulative calibration errors in the positioning of the stepper motor. The design files are freely available at the OSF repository [22].

**Figure 2:**
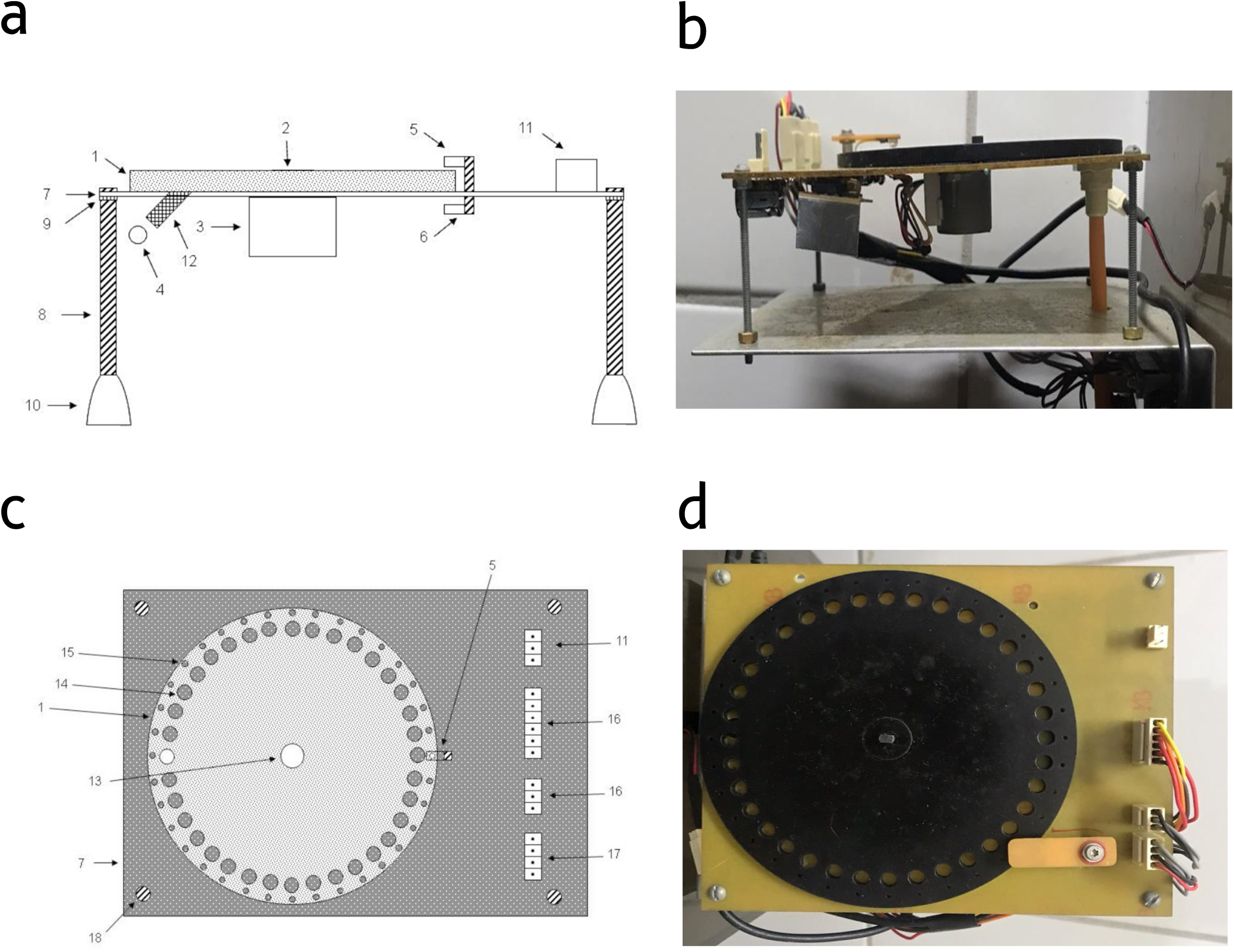
Food pellet dispenser. **(a) Lateral scheme view**, consisting of: pellet release disk (1); stepper motor connector (2); stepper motor (3); food pellet (4); infrared light emitting light (5); infrared light receiver (6); support base (7); threaded support rods (8); nut (9); rod bases (10); circuit power connectors (11); delivery cannula (12). **(b) Lateral view** of the food pellet dispenser. **(c) Top scheme view** Consisting of: pellet release disk (1); infrared light-emitting diode (5); support base (7); circuit power connectors (11); stepper motor connector (13); larger hole (14); smaller hole (15); circuit connectors (16); circuit connectors (17); screw (18) **(d) Top view** of the food pellet dispenser.

### 5.2 Controller

We set up an auxiliary electronic circuit (Fig. 3) to power the Arduino, generate the visual stimuli, monitor the lever and IR sensor, and run the stepper motor coils. The print circuit board (PCB) design files are freely available at the OSF repository [22].

**Figure 3:**
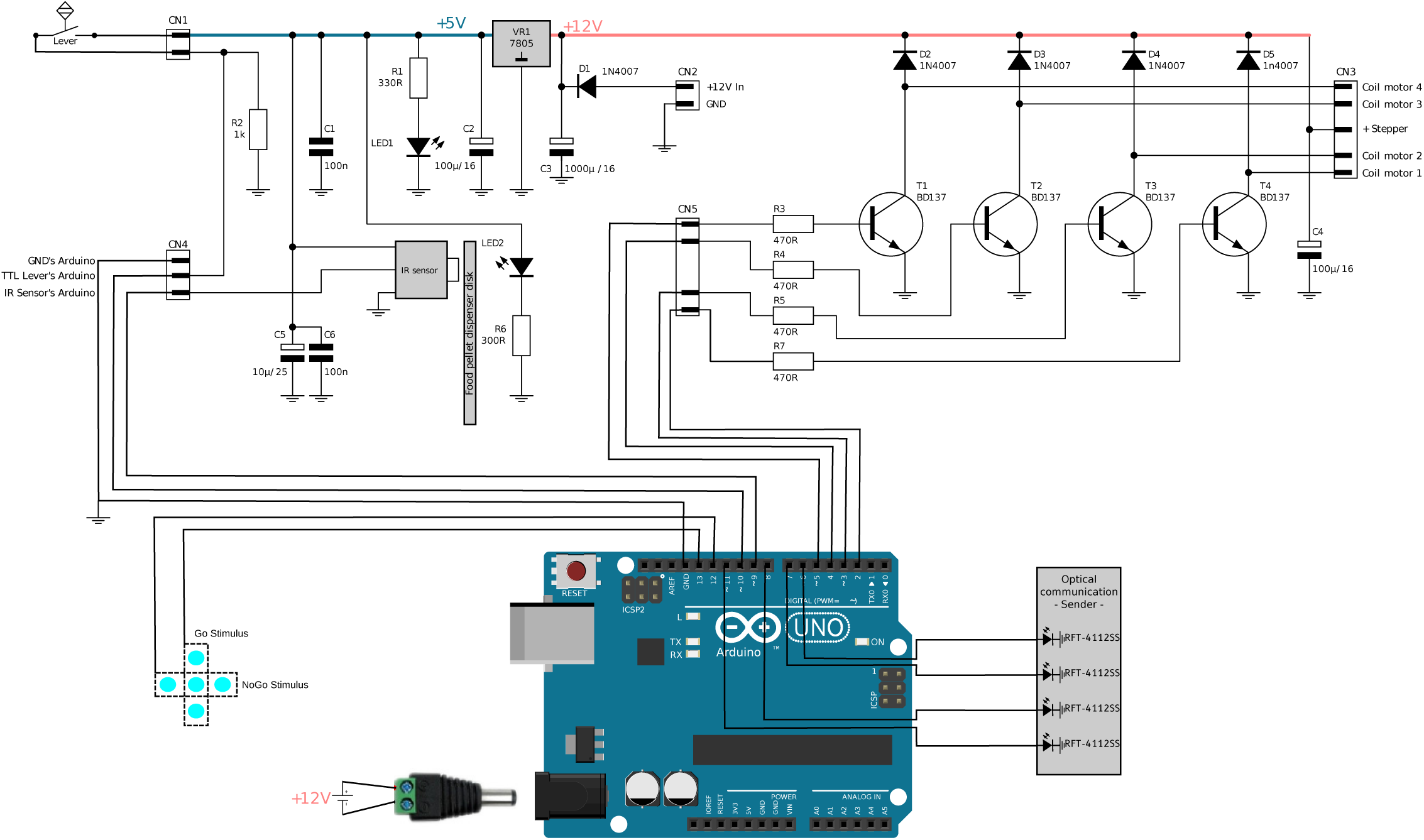
Circuit diagram of auxiliary electronic. Arduino Uno sends and receives TTL controlling signals. Based on the +12V battery, a voltage regulator (VR 7805) is the supplier of two power lines for the electronic components: +5V and +12V. The +5V (in blue) supplies to two LEDs, an infrared receiver (TSOP 1838), and the TTL level to the switch (CN1) used on the lever inside the box. The LED1 is the power-on light, and the LED2 is the emitter to the infrared sensor (IR sensor) under the Food pellet dispenser disk. The +12V (in light red) is the supplier for sequence to the transistors (T1–T4, BD137), which mediate the controlling signals to stepper motor (CN3) from output digital ports in Arduino. Arduino digital ports also send TTL signal for optical communication and were converted to light by RFT-4112SS modules.

Such as is shown in Fig. 3, the auxiliary electronic circuit works with two power supplies: +12 V and +5 V. The +12 V line powers the Arduino (Arduino Uno) and the stepper motor (24BYJ48, with four coils). Each motor coil connects to the collector pin of a PNP bipolar junction transistor (BD137, Fig. 3). Each transistor, at the collector pin, provides a +12 V TTL signal, controlled by a +5 V TTL signal in its base, from an Arduino I/O pin.

The +5 V line was created based on voltage regulator 7805 and powers the infrared sensor and the circuit responsible for detecting lever presses – sending a TTL to the Arduino board when the lever is pressed – and presenting visual stimuli.

The Arduino board also produces two types of actions: it presents the visual stimuli, and it activates the stepper motor. In both functions, the board uses its digital I/O ports. Ports 12 and 13 present one visual stimulus each, each consisting of an illuminated pattern of LEDs. So ports 12 and 13 only turn on/off the respective pattern. Ports 3,4,5,6 control the stepper motor. Upon the moment of the reward, the Arduino sequentially activates the stepper motor rotating the dispenser disk until it releases a sugar pellet in the conduit, which takes the pellet to the cubicle by gravity. An infrared light sensor (Tsop1838) detects the rotation of the disk, through the small hole, and sends a TTL signal to the port 11 Arduino digital port, which in turn stops the disk rotation.

### 5.3 Optical communication

Our system can record the behavioral activity in two different ways: through the serial port of a computer connected to the Arduino and through the TTL signals generated by the Arduino for the acquisition system (Figure 4).

**Figure 4:**
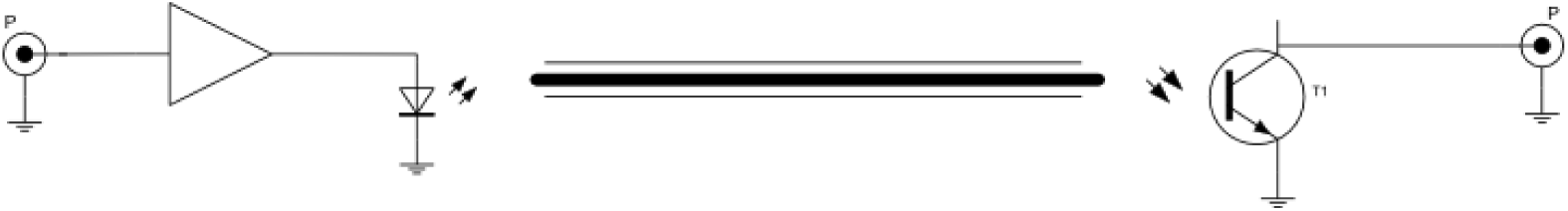
Optical communication scheme. A TTL signal comes out of a digital Arduino port is amplified and converted into light through a LED. The light is transmitted through an optical fiber to the terminal where it will be converted into a TTL signal.

The serial port is helpful when recording behavioral data exclusively – with no need to synchronize with the electrophysiology equipment. We defined a code to each behavioral event, combining the time stamp and the event code. For example, if the rat pressed the lever – code 001 in our case – 823 ms after the session beginning, the serial port of the Arduino sends the following string “823.001”. Such a method encodes the information from all events and their time of occurrence during the analysis.

When recording electrophysiological data along with behavior, we inactivated the Arduino’s serial for recording behavioral activity. The objective was to reduce unnecessary sources of electrical noise that the computer could insert into the system. Instead, our implementation employed an optical fiber to send the synchronization pulses from the Arduino to the Tucker Davis Technologies RZ2 acquisition system. This same arrangement works with the OpenEphys system and may work with virtually any other electrophysiology recording system equipped with TTL input. The TTL signal from the Arduino was converted to a red light pulse by a fiber optic module (RFT-4112SS). In a second step, a phototransistor converts the light signal back to TTL pulses that proceed to the electrophysiology recording system. The TDT recording system has 8 ADCs in its front panel that acquire the synchronization pulses in parallel with the electrophysiological data.

## 6. Operation instructions

The Arduino has an integrated development environment (IDE). The software was written in a C language and loaded directly into the Arduino board. Here we exemplify with three behavioral protocols we implemented and tested.

### 6.1 Fixed Ration sessions

Before initiating the experimental protocols of Differential Reinforcement of Response Duration (DRRD) and Simple reaction-time (SRT) the rat performed a fixed-ratio 1 (FR1) protocol. In these sessions the animal learned to associate the lever with the delivery of a sugar pellet (50 mg sucrose), which was used as reinforcement. The visual stimulus was kept on for the entire session (60 minutes). The trial began when the animal pressed the lever. Only when the lever was released – and irrespective of the duration of the lever press – the Arduino board delivered a sugar pellet. Animals were considered trained when they received 100 sugar pellets within a single one hour session. During the FR1 sessions, the Arduino board monitors a digital port that receives information from the lever, stores it and generates TTL signal relaying the state of the lever. It also sends out a signal that rotates of the disk (carousel) that delivers the sugar pellets.

### 6.2 Differential Reinforcement of Response Duration

The differential reinforcement of response duration (DRRD) task requires that the rat sustains a behavioral response for a minimum amount of time (criterion) [23, 24]. The trial was initiated when the animal pressed the lever. The animals should depress the levers down for more than 1500 ms to receive the reward. When the animal releases the lever before this minimum time, the response is considered premature and is not reinforced.

During DRRD sessions, for each lever press, the Arduino board monitored for how long the rat kept the lever pressed. If it kept the bar pressed for a period longer than the criteria, the Arduino released the reward initiating the stepper motor that rotates the carousel until a sugar pellet drops. More details about the this task can be found in Reyes et al. [24] under the name of “Fixed” protocol. We used three TTL signals to encode the following events in the SRT experiments: 1) Bar state; 2) Food delivery/Success trial; 3: Fail trial.

### 6.3 Simple reaction-time

The Simple reaction-time (SRT) task requires that the animal hold the lever pressed for a minimum time to characterize engagement (500 ms). When this waiting period is over, the system presents a stimulus preceded by a delay – randomly presented and uniformly distributed from 0 to 200 ms.

The animal receives a reward if it releases the lever within a period of reaction time (300 ms in this example) after the stimulus presentation (reinforced responses). Responses are not reinforced if the rat releases the bar before the visual stimulus (premature trial) or if the rat releases the bar after the reaction time (Fail trial).

In addition to monitoring the lever status and the reward, this software also controls the stimulus presentation. We used a modified version of the protocol used by Narayanan and coleagues [25].

We used four TTL signals to encode the following events in the SRT experiments: 1) Bar state; 2) Stimulus on/off; 3) Food delivery/Success trial; 4) Fail trial.

### 6.4 GO-NOGO task

We have also implemented a GO-NOGO task. This task also requires the animal to hold the lever down for a minimum time to characterize engagement (500 ms). After this waiting period, the visual stimulus is presented (randomly sampled between GO and NOGO stimuli). For the visual stimulus, we used arrays of either vertical or horizontal LED patterns, associated with GO and NOGO trials, respectively. If the stimulus conveys a GO trials, animals must release the lever within 500 ms after the stimulus onset to be rewarded with a sugar pellet. If the stimulus indicates a NOGO trial, the animal must hold the lever for more than 500 ms after the stimulus onset to be rewarded. If the rat releases the bar before the visual stimulus, the trial is not rewarded (premature trial).

We used five TTL signals to code the events in the SRT experiments: 1) Bar state; 2) stimulus GO on/off; 3) Stimulus NOGO on/off 4) Food delivery/Success trial; 5) Failed trial.

## 7. Validation and characterization

### 7.1 Subjects

For the experiments described here, we used six rat Long-Evans *Rattus norvegicus*. They were obtained from the animal house from the Laboratory of Computational and Systems Neuroscience, Department of Physics, Universidade Federal de Pernambuco (Recife, Brazil). They weighed 300 g - 370 g and thew were 12 to 15 weeks old. The rats were housed in cages and maintained in the light / dark cycle of 12 h and had free access to water and commercial feed Presence (Neovia - Paulínica, Brazil) until the beginning of the training. The animals were kept under food restriction to keep them at 90 % of the *ad libitum* weight. The experimental protocol was approved by the Ethics Committee on Animal Use (CEUA) of UFPE (CEUA: 24/2016), in accordance with the basic principles for research animals established by the National Council for the Control of Animal Experimentation (CONCEA). The rats were divided into two groups according to the behavioral task they were trained: *SRT* (n=3) *DRRD* (n=3).

### 7.2 Surgical procedures and Recordings

The microwire arrays were built with 50 *µm* tungsten wires, coated with Teflon (California Fine Wire Company), soldered in a printed circuit board, and connected to a miniature connector (Omnetics Connector Corporation, Minneapolis, MN). The microwire arrays were designed for five different cortical regions: PFC, S1, M1, PPC and V1. Table 1 shows the target stereotaxic coordinates of each region according to [26]. The arrays were placed in rats under deep ketamine (150 mg/kg i.p.) and xylazine (5 mg/kg i.p.) anesthesia, using a stereotaxic head holder. through a surgical procedure described by Wiest et al. [27]. Rats were allowed seven days to recover from the surgery and start the behavioral tasks.

**Table 1:** Steriotactic coordinates and arrangement of arrays used in the DRRD task according to [26].

|  | **PFC** | **M1** | **S1** | **PPC** | **V1** |
| --- | --- | --- | --- | --- | --- |
| AP (mm) | +4.00 | 2.00 | 0.84 | -4.36 | -7.20 |
| ML (mm) | 0.50 | 2.50 | 5.00 | 4.20 | 3.50 |
| DV (mm) | -3.70 | -1.50 | -2.00 | -1.80 | -1.50 |

Electrophysiological data were recorded simultaneously from five regions, along with the behavioral activity, at a sampling rate of 24414 Hz, and with a bandwidth of 0.1 to 8 kHz. A ZIF-Clip*®* connector attached the rats to the PZ2 aplifier via a commutator (AC32, Tucker-Davis Technologies, Alachua, FL), which allows the rats to freely move during the recordings. Besides, PZ2 amplifier system (TDT) was responsible to record the local field potential (LFP). Behavioral activity was recorded as digital inputs together with the electrophysiology.

### 7.3 Data analysis

We used the Matlab toolbox Wave Clus [28] to do spike sorting in the raw data. Both off-line raw extracellular recordings and behavioral data were stored locally and in the cloud [29] for later processing, including automatic and manual spike sorting.

To evaluate LFP noise we calculated the spectrogram and coherence matrix. The short-time spectral analysis around the time *t* was calculated by using the short-time Fourier transform (SFFT). Thus, we used the Eq. 1, to calculate the spectrogram, *S*_*i*_(*t, ω*), of *i*-th LFP channel, *x*_*i*_ [30, 31].

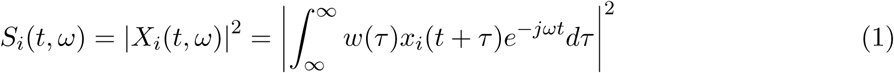

where *w*(*τ*) is a *T* -width time window. Given a pair of time-series of LFP channels, *x*_*i*_(*t*) and *x*_*j*_(*t*) the coherence between them is given by the Eq. 2:

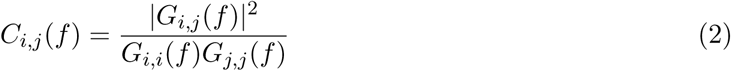

where, *G*_*i,j*_ (*f*) is the cross-spectral density between those time-series *x*_*i*_ and *x*_*j*_; whereas *G*_*k,k*_ is the auto-spectral density of the *k*-th time-series [30]. The element *a*_*i,j*_ in the coherence matrix, such as is shown in Fig 6c, was calculated as the average coherence in a frequency range above the low frequencies of the LFP, where the spikes power spectral begins (300–500Hz). All LFP and spike data were analysed custom-made Python routines, based on open packages: Numpy (1.16.4), Scipy (1.3.0) and Matplotlib (3.1.0), Pandas (0.25.0) and Seaborn (0.9.0). We calculated the response distribution by convolving the histogram of the response duration in 200-ms bins with a Gaussian kernel.

**Figure 5:**
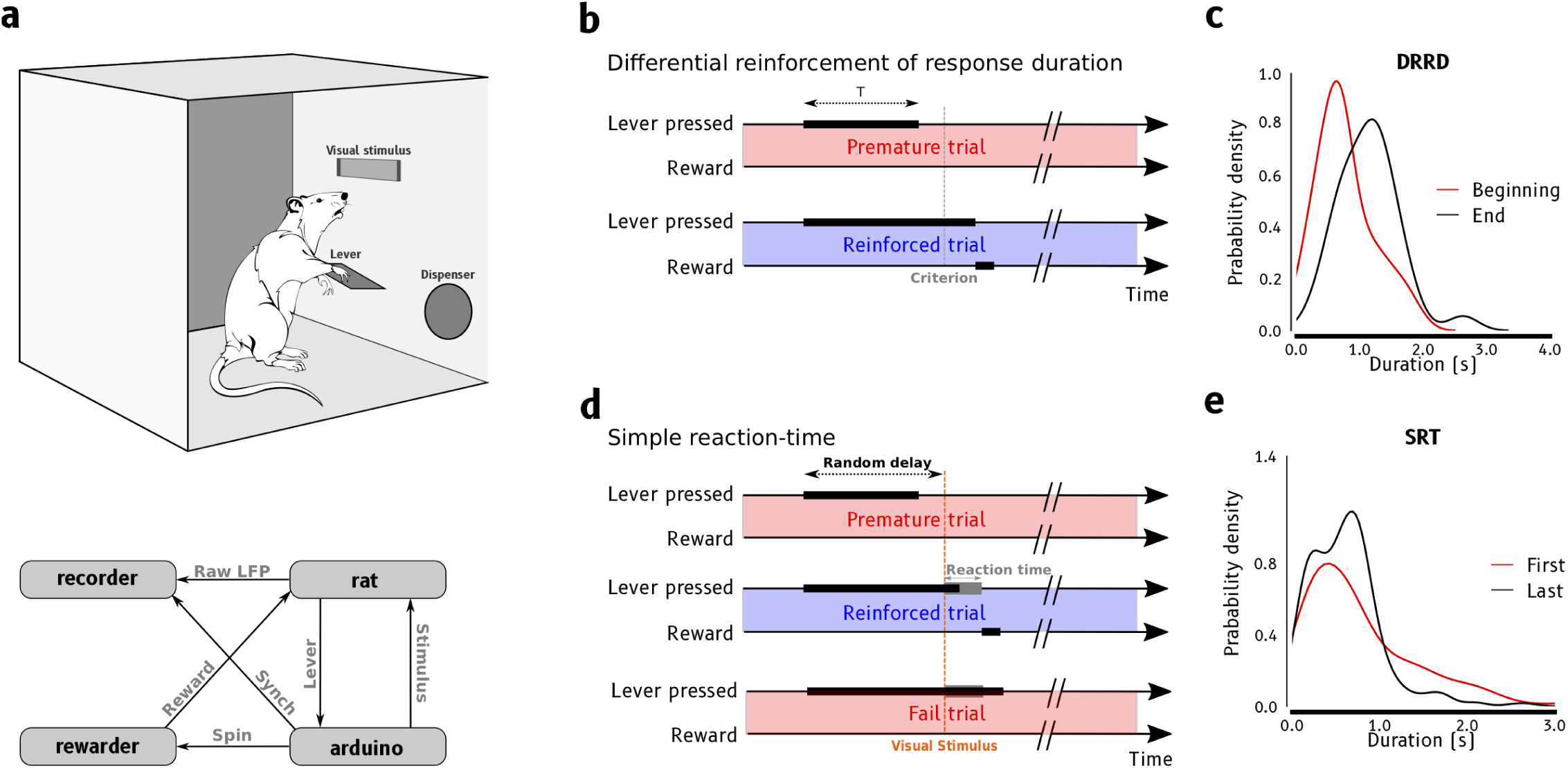
Overview of the experimental behavioral box. (**a**) **top**: operant conditioning chamber containing the LED (for providing visual stimuli), lever, pellet dispenser, and underlying electronic components. **bottom**: Diagram of main components, and its interactions, in the current proposal: the recording system (recorder), which records both behavioral and wideband LFP synchronized data; the animal (rat); the reward device (dispenser), which is a pellet dispenser, controlled by the microcontroller board (arduino) delivers the reward to animal; and the microcontroller board (arduino), a Arduino Uno that controls the reward device and informs the animal behavior along the time. (**b**) Description for differential reinforcement of response duration (DRRD) behavioral tasks, for more details see in the section 6.2. (**c**) Probability density functions of response duration at the beginning (red) and the end (black) in the seventh DRRD session. (**d**) Description for simple reaction-time (SRT) behavioral tasks, for more details see in the section 6.3. (**e**) Probability density functions of response duration at the first (red) and at the last (black) of the SRT behavioral tasks.

**Figure 6:**
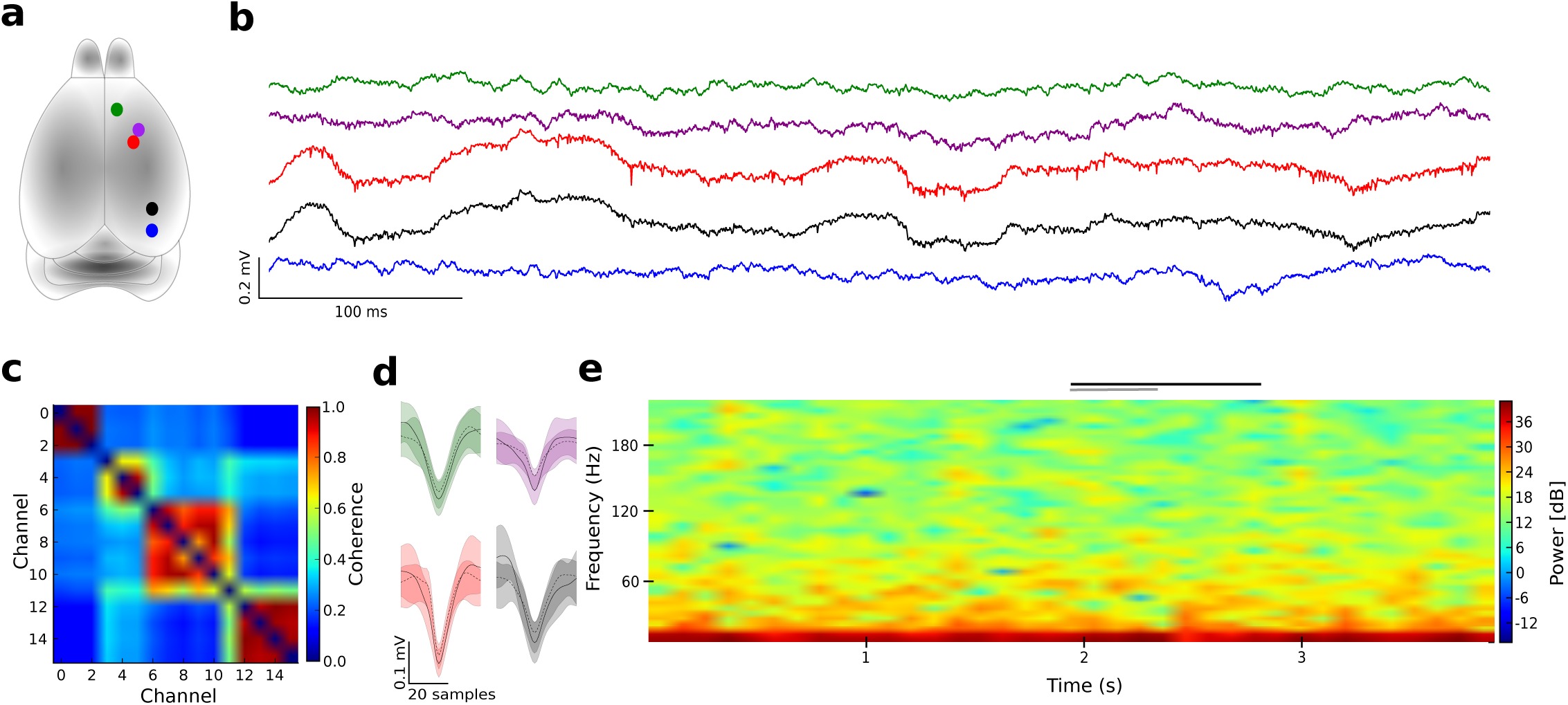
Samples of electrophysiological recordings in a behaving animal. (**a**) Whole rat brain using a color code along the implanted areas: prefrontal cortex (PFC, green), primary motor cortex (M1, purple), primary somatosensory cortex (S1, red), posterior parietal cortex (PPC, black), and primary visual cortex (V1, blue). (**b**) Sample of simultaneous raw wideband LFP signals, according the color code in (a), while the rat performed FR1 task. (**c**) Coherence matrix between the raw LFP across channels, at low frequencies (0-50 Hz). (**d**) Samples of two spike waveforms found in four areas, using color shades based on the color code in (a); within each the cluster shadow represents the amplitude range (mean *±* standard deviation), and filled lines represents the average amplitude, along the waveform samples. (**e**) Sample of spectrogram for low frequencies (up to 200 Hz) of a raw extracellular potential in the PPC, along 4s; upper horizontal black bar represents the period for which the animal kept the lever pressed, while the nearby gray line represents the period in which the reward device was powered on. Local powerline works at 60 Hz.

### 7.4 Results

We assessed the behavior of rats in the DRRD sessions by measuring the duration of the lever presses – the time elapsed between the lever press and the lever release. Since animals produce a high number of lever presses in a session, we characterize the behavior in terms of the probability density of the lever press duration. These curves represent the probability as a function of the time (duration in seconds). In DRRD sessions, animals must press the lever for more than 1.5 s to get rewarded. During FR1 training, the average duration was around 0.5 s. Hence, during training, we expect the distribution of responses to shift rightward, representing an increase in the mean response duration. There was a significant difference between the distribution of response duration at the beginning and the end of the DRRD training in a single session (Fig. 5c, *p <* 0.001, Mann-Whitney). Where we found larger means for response duration during the end sessions (1.14 s) when compared with the beginning session (0.77 s). The proportion of short responses decreases over responses around the criterion (1.5 s). Our data show that a small number of training days elicited a significant change in behavior, corroborating with Reyes et al. [24].

We found similar differences over sessions in the SRT task (Fig. 5e), where responses were on average longer in the last session (0.81 second) when compared with the first sessions (0.61 second; *p* = 0.020, Mann-Whitney). These results show that rats learned the task because the proportion of premature responses decreases while the proportion of responses after the stimulus (SRT trials) increases. After the training sessions, the peak of the distribution of response times increases and is compatible with the expected correct response time. Our data show that the training elicited a significant change in behavior along with the sessions in both behavioral tasks.

An essential advantage of the OCC presented here is that it does not induce noise in the electrophysiological recordings. In Fig. 6 we show the raw signals (recorded from PFC, M1, S1, PPC and V1 cortical areas) during an FR1 task (panel b) and the spikes extracted from the raw data (through a spike sorting process) confirming the quality of the recordings (panel d). Such very low voltage (few hundreds of micro-volts) recordings are quite sensitive to electromagnetic artifacts, and two events are critical: the lever press (black line) and the activation of the stepper motor (gray line). If the electric circuits are not properly isolated, such events typically create electrical artifacts in the recordings due to static energy discharge when the animal touches the lever, sudden changes in potential upon lever press (switches), or to inductive charges from the stepper motor coils. Our recording remained unaltered by these events shown in panel (Fig. 6b). The spectrograms of the LFP signals were also free of artifacts (Fig. 6e) – no increase in the fundamental (60 Hz), harmonics (120 and 180 Hz) levels, nor in higher frequencies were observed.

The coherence map (Fig. 6c) also tests the quality in the recordings. It is possible to identify four broad clusters of high coherence (warmer colors). It is expected that the coherence was high for electrodes in the same cortical region, but low for electrodes more distant apart. The first cluster corresponds to the electrodes inserted in PFC, the second in M1, the third in S1 and PPC (highly synchronized) and the fourth in V1, confirming that the high coherence was only observed in adjacent electrodes.

## 9. Acknowledgements

This project was partially supported by CNPq (Grant Project No. 430993/2016-1), FACEPE (Grants APQ 0826-1.05/15 and APQ 0642-1.05/18), CAPES (L.A.A.A; N.A.P.V. Grant No. 88887.131435/2016- and FAPESP (Grant 2018/20277-0; G.C.T Grant 2016/18914-7). N.A.P.V. thanks Dr. Sidarta Ribeiro’s lab by the part of this work supported by the high performance computing facilities of NPAD/UFRN. We thank OSF project to provide the cloud storage for this study.

